# The frequency dependence of EEG-speech envelope tracking does not reflect the rates of speech units or pitch

**DOI:** 10.64898/2026.06.14.732143

**Authors:** Mike Thornton, Tobias Reichenbach

## Abstract

During speech listening, neural activity partly synchronises to fluctuations in the acoustic amplitude envelope of the stimulus. This neural tracking of the speech envelope has frequently been linked to the processing of acoustic features such as syllabic rhythmicity and pitch periodicity. However, it has not yet been investigated to which degree the frequency dependence of neural envelope tracking reflects the rate of acoustic and linguistic speech units. Here, using a large dataset of electroencephalographic (EEG) responses from participants who listened to naturalistic speech, we quantified neural tracking of the speech envelope across a wide range of modulation frequencies using coherence analysis. The coherence profile exhibited distinct peaks in the low-delta band (0.2–2 Hz), the theta–alpha band (4–15 Hz), and at higher frequencies near 45, 95, and 175 Hz. We show that this structure is independent of the rates of phonemes, syllabes and words as well as of pitch periodicity. Using temporal response functions (TRFs), we further show that neural envelope tracking in the gamma frequency band (30–250 Hz) is primarily driven by two clusters of neural generators with latencies of approximately 8 ms and 25 ms, likely located in the rostral brainstem and across the thalamocortical pathway, respectively. Together, these results highlight the interplay of different neural mechanisms and sources in shaping the neural envelope tracking, and lead to important methodological and interpretative considerations for studies that assess neural envelope tracking within narrow frequency bands.

## 1 Introduction

Neural coupling to sound amplitude is a ubiquitous phenomenon in auditory neuroscience. A well-known example from electrophysiology is the auditory steady-state response (ASSR), in which electroencephalography (EEG) signals phase-lock to periodic amplitude modulations of simple stimuli such as sinusoidally modulated noise or tones. Notably, this coupling depends upon the modulation rate, with ASSR amplitude peaking near modulation frequencies of 40 Hz (Galambos et al., 1981).

Recently, methods from system identification have been adopted to assess the synchrony between neural activity and speech features – including the acoustic amplitude envelope – during the perception of continuous spoken narratives (Crosse et al., 2016; Lalor et al., 2009). In this framework, encoding and decoding models are used to relate neural time series to speech features such as the amplitude modulations of the speech stimulus, typically within a predefined frequency band. Model performance is then quantified using a goodness-of-fit metric such as the Pearson correlation coefficient, which is commonly interpreted as a measure of neural speech tracking (Crosse et al., 2016).

The application of these methods has, over the past years, uncovered associations between the neural tracking of the speech envelope and cognitive aspects of speech processing, including selective attention and speech-in-noise comprehension (Ding and Simon, 2012; Etard and Reichenbach, 2019; O’Sullivan et al., 2014; Vanthornhout et al., 2018). In parallel, the convergence of several observations has led to the suggestion that neural envelope tracking may reflect a mechanism for segmenting syllables in connected speech, particularly within the theta (4-8 Hz) range. First, the long-term modulation spectrum of continuous speech exhibits prominent energy at the syllabic rate, with sharp rising edges in the time domain reliably indicating the onset of syllabic nuclei (Oganian and Chang, 2019; Rosen, 1992; Zhang et al., 2023). Second, neural envelope tracking is thought to be particularly strong near the syllabic rate (Issa et al., 2024; Peelle et al., 2012). Third, some behavioural studies indicate the existence of a perceptual primitive of speech with a “syllable-sized” timescale of approximately 200 ms (Arai and Greenberg, 1998; Saberi and Perrott, 1999). Some authors have employed similar arguments to further suggest a link between delta-band (1-4 Hz) neural envelope tracking and the prosodic rhythm of continuous speech, and between low-gamma (30-50 Hz) activity and the timescales of phonetic transitions and manner cues (Bourguignon et al., 2012; Giraud and Poeppel, 2012; Meyer, 2017; Rosen, 1992).

Motivated by these accounts, many neuroimaging studies assessed neural envelope tracking within narrow frequency bands selected to align with the temporal rates of putative speech units. Studies employing such an approach reported, for instance, a functional dissociation between frequency bands, with delta-band tracking appearing more strongly modulated by cognitive factors, and theta-band tracking more closely tied to the acoustic conditions of speech presentation (Etard and Reichenbach, 2019; Keitel et al., 2018; Molinaro and Lizarazu, 2018). However, a fine-grained characterisation of neural speech-envelope tracking across modulation frequencies is still lacking.

Neural envelope tracking of continuous speech can also be observed at substantially higher frequencies, including those comparable to the fundamental frequency of speech (F0) (Hertrich et al., 2011; Kulasingham et al., 2020). Such responses have attracted particular interest, since they are modulated both by selective attention to speech (Commuri et al., 2023; Schüller et al., 2023) and by lexical word frequency (Kegler et al., 2022). However, it remains unclear to what extent these responses reflect neural tracking of pitch periodicity. Moreover, the modulation-frequency dependence of neural speech-envelope tracking across the broader gamma band has not yet been systematically characterised.

The anatomical origins of the neural envelope tracking in the gamma band remain an active area of investigation. Evidence from EEG studies indicated that early components of the response are subcortical with a main contribution from the rostral brainstem. This interpretation was supported by the similarity of the response latency to that of wave V of click-evoked brainstem response and through computational modelling (Bachmann et al., 2021; Maddox & Lee, 2018; Saiz-Aĺıa & Reichenbach, 2020). In contrast, MEG studies revealed an additional later component with a focal origin in the auditory cortex and with a significant right-hemisphere bias (Kulasingham et al., 2020; Schüller et al., 2024). Polonenko and Maddox (2021) further demonstrated that EEG responses in the gamma range include an intermediate, middle-latency component that emerges when envelope tracking is assessed across a broader range of modulation frequencies above 30 Hz. Gamma-band envelope tracking therefore reflects neural processing across the span of the auditory pathway. Further characterising its modulation-rate dependence, acoustic specificity, and anatomical origins could provide valuable insight into the input and organisation of central auditory processing, with potential clinical applications.

In this work, we sought to characterise how neural envelope tracking varies across a wide range of modulation frequencies. To this end, we assessed spectral coherence between the speech envelope and EEG signals recorded from a large cohort of participants who listened to naturalistic speech in the form of audiobooks and podcasts. Owing to the size of the dataset and its high sampling rate of 1,024 Hz, spectral coherence could be estimated with a particularly high frequency resolution. We hypothesized that, due to the role of the neural tracking in speech processing, the coherence spectrum might exhibit peaks at frequencies corresponding to the relevant speech rhythms or periodicities. Conversely, the absence of such speech-rhythm related peaks would indicate other mechanisms that shape the frequency dependence of the neural speech tracking, such as interference between the far-field contributions of distinct neural generators along the auditory pathway (Tichko and Skoe, 2017).

## 2 Methods

### 2.1 Dataset

We analysed data from the SparrKULee dataset, a large, publicly-available EEG dataset measured from participants who listened to naturalistic speech in the form of audiobooks and podcasts (Accou et al., 2024). Data from 81 participants were made available and are included in the present study. Participants were young (18–30 years; mean age 21.5 years), native Flemish speakers with normal pure-tone audiometric thresholds. The dataset is imbalanced with respect to sex (10 males, 71 females). Sixty-nine participants were right-handed. On average, each participant listened to two hours of continuous speech material, segmented into trials of approximately 15 minutes. Between trials, participants answered comprehension questions to encourage sustained attention to the stimuli.

Electroencephalography (EEG) was recorded from 64 scalp electrodes using a BioSemi ActiveTwo system with a BioSemi 64-channel active electrode cap (BioSemi, Amsterdam, The Netherlands). Signals were acquired at a sampling rate of 8,192 Hz. However, the publicly-available data were shared at a reduced sampling rate of 1,024 Hz. During stimulus presentation, an auxiliary audio channel containing regularly spaced trigger pulses was recorded by the ActiveTwo system. These pulses were used to correct for temporal drift between the amplifier clock and the stimulus presentation sound card, enabling precise temporal alignment of the speech stimulus and the EEG recordings. The EEG signals were re-referenced to the average of the 64 scalp channels prior to data analysis.

Auditory stimuli were presented diotically (identical signals to both ears) via insert earphones (Etymotic ER-3A, Etymotic Research, Elk Grove Village, IL, USA). The speech material comprised a variety of audiobooks and podcasts. All participants listened to a common reference stimulus (“audiobook 1”), whereas the remaining stimuli varied across participants. An overview of the prevalence of each audiobook and podcast within the dataset is shown in Figure 1a. All speech material was presented in Flemish Dutch and provided a range of talkers with different fundamental frequencies and vocal characteristics.

**Figure 1:**
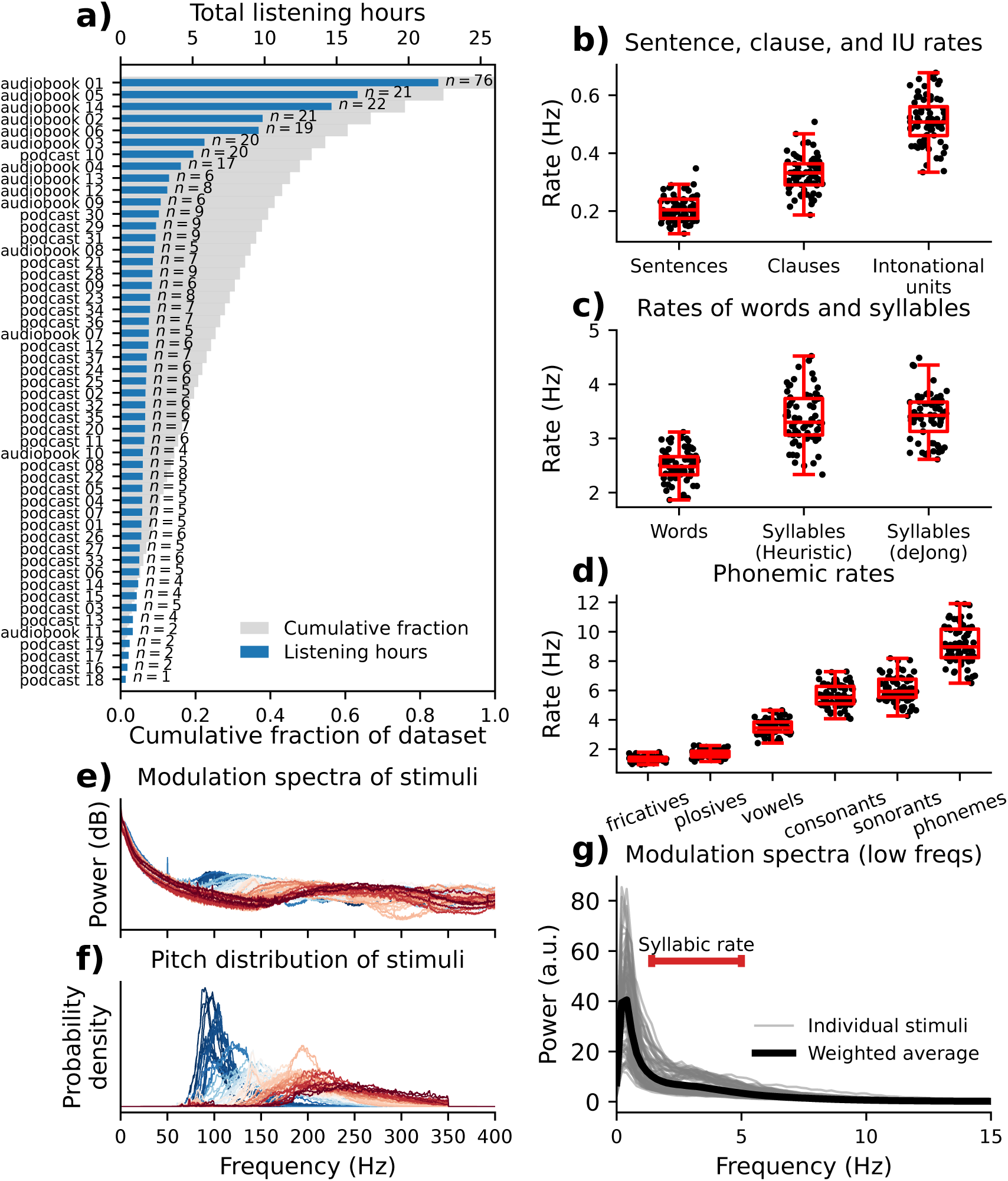
Characteristics of the speech stimuli in the SparrKULee dataset. **a)** Distribution of stimulus exposure across participants. The number of listeners (*n*), total listening time (*n* multiplied by stimulus duration), and cumulative contributions to the dataset are shown. **b-d)** Average rates of selected speech units. Rates are defined as the number of units per second; each data point corresponds to a single stimulus. **e)** Modulation spectra (envelope power spectral density) of the stimuli. Thin coloured lines show individual stimuli, with colours varying with median pitch. Blue denotes a low median pitch around 100 Hz, and red a high median pitch above 200 Hz. **f)** Pitch distributions for individual stimuli, with colours again indicating median pitch as in panel e. **g)** Low-frequency (0–15 Hz) modulation spectra. Many stimuli exhibit an elbow around 2.5 Hz, reflecting syllabic rhythmicity in the 2-5 Hz range (indicated in red). The thick black line shows the average spectrum, weighted by stimulus prevalence (panel a). coherence patterns. Coherence near both 45 Hz and 94 Hz is strongest near the frontal electrode Fz, whereas coherence near the highest-frequency peak at 173 Hz is more centrally clustered along the midline. Above 250 Hz, the coherence spectrum descends into the noise level.

### 2.2 Envelope extraction

The acoustic amplitude envelope of the speech signal was extracted from the raw audio (sampled at 48 kHz) as follows. First, a low-pass filter was applied to the audio signal, in order to mimic the frequency response of the ER-3A insert earphones (low-pass Remez filter of order 19 with a passband edge of 3,500 Hz, a transition bandwidth of 5,500 Hz, and a stop-band attenuation of 45 dB). The envelope was derived from the filtered audio using a coarse model of the auditory periphery based on a gammatone filterbank, following the approach of Biesmans et al. (2017). The audio was passed through a bank of 50 gammatone filters spaced evenly on an equivalent rectangular bandwidth (ERB) scale between 50 Hz and 8 kHz. The gammatone filterbank was implemented in brian2hears (version 2.9.0) using default parameter settings (Fontaine et al., 2011). The output of each sub-band was half-wave rectified and subjected to power-law compression with an exponent of 0.6. The resulting sub-band signals were resampled to 1,024 Hz using `scipy.signal.resample_poly` (SciPy version 1.15.3, Virtanen et al. (2020)), which applies a linear-phase FIR anti-aliasing low-pass filter (Kaiser window with *β* = 5.0, a cutoff frequency of 512 Hz and a transition bandwidth of 0.39 Hz. The stopband attenuation was 60 dB). The broadband envelope was obtained by averaging across sub-bands of the resulting cochleogram.

### 2.3 Annotation of speech units

Because neural envelope tracking is often linked to prosodic structure in the low-delta range (∼ 0.5 Hz), syllable-level processing in the theta band (4–8 Hz), and pitch periodicity in the high-gamma range (on the order of 100 Hz), we extracted representations of these features and compared their rates to the peak frequencies in the EEG-speech envelope coherence spectrum.

First, transcripts of the speech material were automatically generated using Whisper, a state-of-the-art speech-to-text model (Radford et al., 2023). We used the implementation available in the WhisperX Python library (Bain et al., 2023, version 3.7.4), employing a model pre-trained on a large corpus of multilingual data (`Hugging Face model card: openai/whisper-large-v3`). The automatically generated transcripts were visually inspected. Textual tokens that differed from their spoken realizations (e.g., numerals, dates, or symbols) were manually converted to their spoken forms. Forced alignment between transcripts and audio was performed using the Montreal Forced Aligner (McAuliffe et al., 2017, version 3.3.8) with its `built-in dutch_cv` acoustic model and pronunciation dictionary (Ahn and Chodroff, 2022). Alignment was conducted on audio segments of 30 s in duration to minimize the propagation of local alignment errors throughout the long (∼ 15 min) recordings. The alignment procedure yielded word-and categorical phoneme-level annotations with onset and offset timestamps.

To characterise prosody, we estimated the onsets of intonation units (IUs) using the algorithm proposed by Inbar et al. (2025). Intonation units are prosodically-defined stretches of speech marked by acoustic boundary cues such as pitch reset, final lengthening, and amplitude modulation. They typically recur on a timescale of approximately 1–2 seconds and contribute to the low-frequency rhythmic structure of speech (Inbar et al., 2025). We note that alternative measures of prosodic structure are possible, and that the relationship between neural envelope tracking—particularly pronounced at this timescale—and prosodic grouping remains at present speculative (see, e.g., Meyer (2017) for an overview). A comprehensive evaluation of other speech units operating at similar timescales (e.g., sentences, clauses, or alternative prosodic measures) is beyond the scope of the present study.

Syllables were identified from the phonemic transcripts. In Dutch, syllables are organized around a vowel-like nucleus, typically flanked by consonant clusters. We therefore defined syllabic nuclei as the temporal midpoints of vowel phonemes, with diphthongs treated as a single nucleus whose timing was defined by the midpoint of the combined diphthong span. To validate this heuristic syllable-identification procedure, as well as the transcription and forced-alignment pipeline, we compared the average syllable rate (total number of syllables divided by audiobook duration) with estimates obtained using an independent acoustic method for detecting syllabic nuclei (de Jong and Wempe, 2009, version three available at: https://sites.google.com/view/uhm-o-meter). The two approaches yielded highly similar results (see Figure 1c).

### 2.4 Estimation of pitch and formant frequency

Instantaneous pitch (F0) was estimated using Praat’s autocorrelation-based pitch-tracking algorithm (Boersma and Weenink, 2021, version 6.1.38). Praat was accessed via the Parselmouth interface for Python (Jadoul et al., 2018, version 0.4.6). Pitch was computed with a frame length of 10 ms, a floor of 50 Hz, and a ceiling of 350 Hz. Unvoiced frames (for which Praat returns *F* 0 = 0) were excluded from subsequent analyses. For subsequent analysis, the instantaneous pitch estimates were binned into histograms with a 0.2 Hz bin width. Figure 1f depicts these histograms, and shows that they align with gamma-band peaks in the long-term modulation spectra of the speech material.

Prior work has suggested that neural envelope tracking may be enhanced in carrier bands near formant frequencies, particularly at modulation rates close to the fundamental frequency of speech (Hertrich et al., 2011). To test this hypothesis, time series of the first three formant frequencies were estimated using Burg’s algorithm as implemented in Praat (frame length = 25 ms, overlap = 10 ms, maximum formant frequency = 5.5 kHz, pre-emphasis = 50 Hz). Because spectral coherence reflects long-term synchrony between signals, each formant time series was summarised as a histogram, whose centre and width characterise the typical range of each formant for a given talker and can be directly compared with spectral coherence as a function of cochleogram centre frequency.

### 2.5 Coherence analysis

The complex spectral coherence between the speech envelope *x*(*t*) and each EEG channel *y*(*t*) is defined as follows:

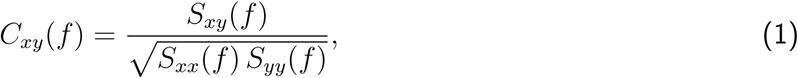

where *S_xy_*(*f*) denotes the cross-spectral density between the envelope and the EEG signal, and *S_xx_*(*f*) and *S_yy_*(*f*) denote the corresponding auto-spectral densities. Spectral densities were estimated using Welch’s method with a window length of 5 s and 80% overlap between successive segments. The 5 s window length reflects a compromise between resolving low-frequency components, which require sufficiently long segments to capture multiple oscillatory cycles, and preserving sensitivity to higher-frequency components, which can vary on much shorter timescales. The resulting frequency resolution was Δ*f* = 1*/T* = 1*/*(5 s) = 0.2 Hz.

The complex coherence was computed separately for each trial and participant. To account for differences in stimulus duration, coherence estimates were combined using a weighted average in which the weights were chosen proportional to the duration of the corresponding audio stimulus (see Figure 1a for an overview). To obtain a single coherence profile across electrodes, the root-mean-square (RMS) of the coherence magnitude was then computed across channels:

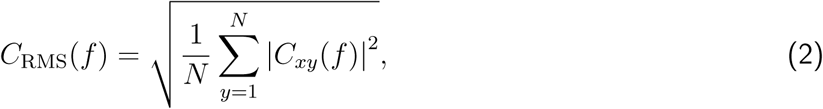

where *N* = 64 is the number of EEG channels.

The resulting coherence spectrum exhibited a number of peaks. We wanted to derive a *p*-value for the statistical significance of each EEG channel’s coherence within a set of frequency bands defined by the widths and centres of these peaks. Therefore, we repeated the complex coherence estimation procedure, but this time using temporally mismatched envelope and EEG segments. The resulting null coherence spectrum exhibited a flat profile with no frequency-specific structure (Figure 2). The complex null coherence was modelled as a circularly symmetric complex normal random variable with independent real and imaginary components. Under this model, the magnitude-squared coherence follows an exponential distribution,

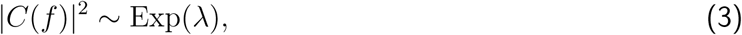

where the rate parameter *λ* is the variance of the underlying complex normal distribution. The parameter *λ* was estimated by fitting an exponential distribution to the empirically observed magnitude-squared noise coherence values. When averaging the magnitude-squared coherence across a given frequency band comprised of *n* adjacent frequency bins, the resulting statistic follows a gamma distribution,

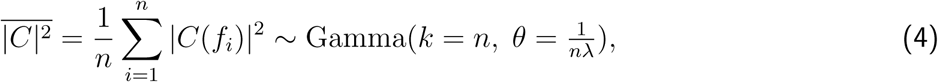

where *k* denotes the shape parameter and *θ* the scale parameter. A single-tailed *p*-value for the mean magnitude-squared coherence within a particular frequency band was then obtained for each EEG channel by evaluating the corresponding percentile of this null distribution. Equation 4 was validated for several values of *n* by visually comparing the theoretical probability density function with histograms obtained from the null coherence profile.

**Figure 2:**
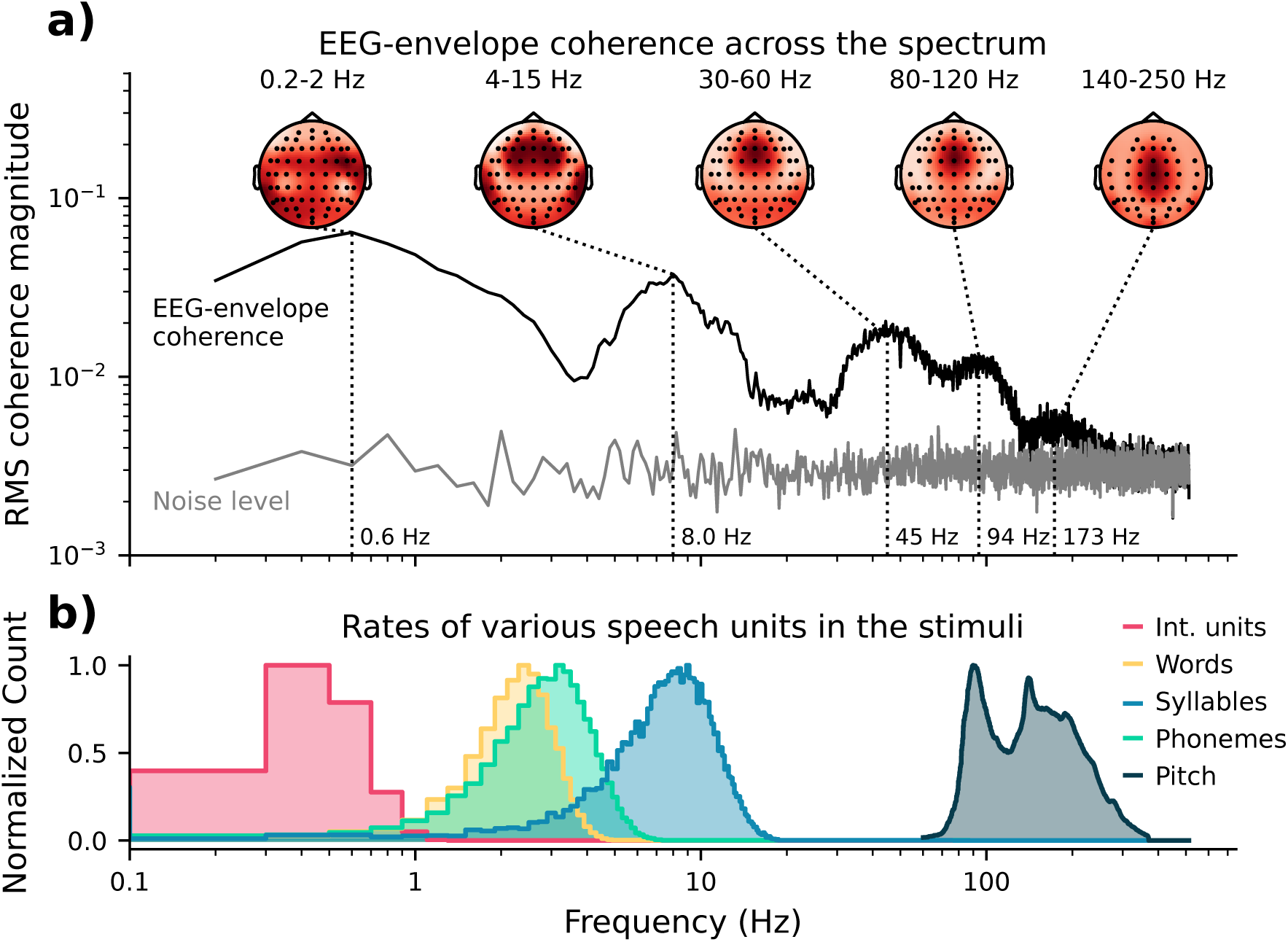
Relation between EEG-speech envelope coherence and the rates of acoustic units. **a)**: Spectral coherence between the EEG signals and the broadband speech envelope, shown on logarithmic axes. The profile reflects the root-mean-square (RMS) coherence magnitude across channels. Topographic maps show the spatial distribution of coherence within frequency bands corresponding to five prominent peaks as well as one shoulder in the spectrum. **b)**: Histograms of the rates of selected speech units and the pitch of the speech material. The pitch distribution is bimodal, reflecting the presence of both male- and female-narrated speech in the dataset. The rates of intonational units, phonemes, and pitch coincide with frequency ranges in which the coherence has maxima. The rates of words and syllables, in contrast, fall into a minimum of the coherence.

### 2.6 Comparison between spectral coherence and the rates of speech units

Section 2.3 described the extraction of timestamps for four speech units: intonation units, words, syllabic nuclei, and phonemes. These timestamps marked unit onsets relative to the start of the audio stimuli. To obtain rates comparable to the coherence spectrum, we counted occurrences of each unit within the same overlapping 5-second windows used for the Welch spectral estimates described above, and divided by window duration to yield rate histograms. Histograms from different auditory stimuli were combined via a weighted-average, with the number of participants who listened to a particular stimulus determining the weight. The peak frequencies and widths of these averaged histograms were then compared to the peaks in the EEG-speech envelope coherence spectrum.

Pitch and formant histograms were also averaged according to stimulus prevalence in the dataset. The resulting pitch distribution can be directly compared to the coherence profile at high modulation frequencies (i.e. in the high-gamma band).

Although we predominantly focused on quantifying the coherence between the EEG measurements and the broadband speech envelope, we also wanted to understand how the envelopes of different acoustic carrier bands might be tracked individually. Therefore, we additionally computed the spectral coherence between the EEG signals and each frequency channel of the gammatone cochleogram.

### 2.7 TRF analysis

While spectral coherence quantifies frequency-specific synchronisation between the speech envelope and EEG signals, it does not provide information on the latencies of the underlying neural responses. We therefore employed temporal response functions (TRFs) to resolve the temporal profile of envelope tracking and to examine its spatial distribution across electrodes as a function of response latency. The TRF analysis was motivated in particular by the triple-peak structure that was observed in the gamma band of the coherence spectrum, which warranted further characterisation in the time domain.

Temporal response functions (TRFs) can be estimated in the frequency domain using ridge regression. Let *x*(*t*) denote the speech envelope and *y_k_*(*t*) the EEG signal at channel *k*. The TRF *h_k_*(*τ*) is defined by a linear time-invariant (LTI) model (Crosse et al., 2016; Lalor et al., 2009):

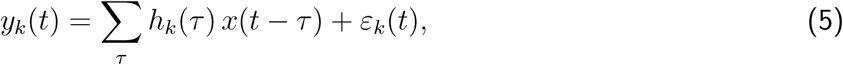

where *ε_k_*(*t*) denotes residual activity not explained by the model. Taking the Fourier transform yields

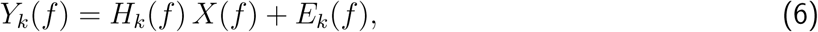

where *H_k_*(*f*) denotes the transfer function. A ridge-regularised frequency-domain estimate of the transfer function is therefore given by

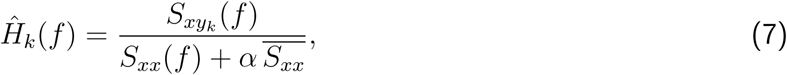

where *S_xyk_* (*f*) and *S_xx_*(*f*) denote the cross- and auto-spectral densities, respectively, and α is a dimensionless regularisation parameter. *S_xx_* and *S_xy_* were taken as the same spectral estimates that were used for the coherence analysis (Welch’s method; 5 s segments; 80% overlap). The regularisation term is scaled by the mean stimulus spectral density *S_xx̅_* across frequency. The parameter α thus quantifies regularisation strength relative to overall stimulus power and results in a frequency-domain formulation equivalent to the time-domain normalisation approach described by Biesmans et al. (2017).

Studies often employ *α* ≈ 1 to improve the stability of TRF estimates. Regularisation is particularly important when training data are limited or when regressors are narrow-band, but it can distort the shape of the estimated TRF, especially for broadband neural signals with frequency-dependent power. In the present work, we used broadband regressors and a large dataset, reducing the need for regularisation; accordingly, we set *α* = 0. Nevertheless, TRFs obtained with *α* = 1 were found to be qualitatively similar to those reported here.

Prior to fitting *H*^^^*_k_*(*f*), the cross-spectral density *S_xy_* (*f*) was normalised by the standard deviation of the EEG signals in the gamma band (30-250 Hz) on a channel-by-channel basis, following a normalisation convention established in several prior EEG studies (Bachmann et al., 2021; Kegler et al., 2022; Van Canneyt et al., 2021). Therefore, the TRFs do not directly reflect the response magnitude in physical dimensions of electric potential. Instead, this normalisation emphasises EEG channels which more faithfully track the regressor signal *x*(*t*) (in this case the speech envelope), resulting in TRFs whose scalp distributions are more readily compared with the spatial distribution of EEG-speech envelope coherence. For comparison, TRFs fitted without this normalisation are shown in Supplementary Figure S1.

The time-domain TRF was obtained via the inverse Fourier transform of *H*^^^*_k_*(*f*):

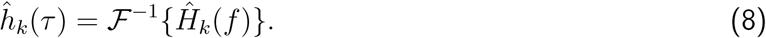

Frequency-specific TRFs were obtained by band-limiting the estimated transfer function prior to inversion. A linear-phase finite impulse response (FIR) band-pass filter was designed using a Kaiser window (60 dB stopband attenuation; transition width 5 Hz; sampling rate 1,024 Hz). The squared magnitude of the filter’s frequency response was applied directly to *H*^^^*_k_*(*f*), yielding a zero-phase band-limited transfer function equivalent to forward–reverse (“filtfilt”) filtering in the time domain. The narrowband TRF was then obtained via the inverse Fourier transform,

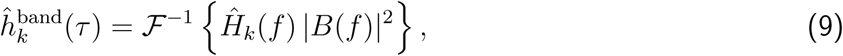

where *B*(*f*) denotes the frequency response of the FIR filter. This filtering approach was also used to remove power line noise at 50 Hz and harmonics thereof (notch filters designed with scipy.signal.iirnotch with a quality factor of 30; harmonics of 50 Hz up to 450 Hz were removed).

Because the TRFs were band-limited in the frequency domain, the resulting time-domain responses necessarily exhibited broad, oscillatory temporal profiles. This reflects the fundamental trade-off between temporal and spectral resolution: restricting a signal to a narrow frequency band inevitably spreads its representation in time (Gabor, 1946). Because such wavelet-like responses are not strictly localized in time, we did not follow the standard approach of analysing the raw TRF waveforms, but rather their instantaneous amplitude:

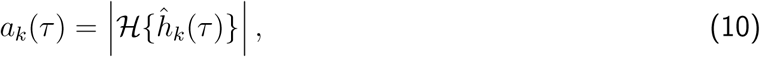

where H{·} denotes the Hilbert transform. The maxima of *a_k_*(*τ*) were interpreted as the temporal loci of envelope-driven neural activity within the selected frequency band. Such an approach has been employed in prior work (Forte et al., 2017; Schüller et al., 2023).

### 2.8 Relationship between the TRF and the spectral coherence

The spectral coherence provides a frequency-domain characterisation of neural envelope tracking that is complementary to the TRFs, because both approaches can be cast in terms of the same linear signal model of equation (5). Substituting the Fourier-domain signal model (equation 6) into the definition for spectral coherence (equation 1) yields the following expression for complex coherence in terms of the transfer function:

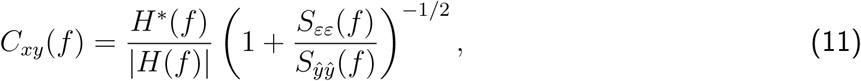

where *S_εε_*(*f*) denotes the auto-spectral density of the residual process *ε*(*t*) defined in equation (5), and *S_y_*_^*y*^_(*f*) = |*H*(*f*)|^2^*S_xx_*(*f*) is the spectral density of the estimated response.

Equation (11) shows that the phase of the complex coherence is determined by the phase of the transfer function, while its magnitude is a number between zero and unity that depends solely on the ratio between the residual spectral density and the spectral density of the estimated response. The squared coherence magnitude therefore reflects the proportion of EEG variance that can be explained by the linear signal model (5) within each frequency bin. It is worth noting that our approach of averaging the *complex* coherence estimates implicitly assumes that the phase relationship between stimulus and response—and therefore the response latency—is consistent across listeners. Substantial inter-participant variability in latency would lead to phase cancellation and attenuated coherence estimates (Keding et al., 2024). This assumption also underpins the common time-domain practice of averaging TRFs across participants.

## 3 Results

### 3.1 Coherence between EEG and the broadband speech envelope

Figure 2a shows the spectral coherence between the EEG signals and the speech envelope. The spectrum exhibits five prominent peaks spanning frequencies from the low-delta to the high-gamma range. In the low-delta band (0.2–2 Hz), coherence reaches a global maximum at 0.6 Hz. The spatial distribution in this range is strongly right-lateralized, with a focal maximum over the right auditory cortex. A second peak occurs at 8 Hz, spanning approximately 4–15 Hz, which is the theta–alpha range. Coherence in the theta-alpha range is distributed symmetrically across the scalp and is consistent with bilateral sources in auditory cortex. In the gamma range, three broader peaks are observed at 45 Hz, 94 Hz, and 173 Hz. Compared to the lower-frequency peaks, these are much wider and show more centrally distributed

#### No relationship between the theta-alpha peak and the rates of speech units

The rates of selected speech features are shown in the lower panel of Figure 2, on the same frequency axis as the coherence plot. The average syllabic rate does not align with any of the peaks in the coherence, but instead falls near the local minimum between the low-delta and the theta–alpha peaks at around 3.5 Hz. Nor does the word rate align with any of the peaks in the coherence spectrum. In contrast, a correspondence can be observed between the phonemic rate and the theta–alpha peak. However, this alignment appears to be coincidental: Figure 3a shows that when the coherence analysis is repeated for subgroups of stimuli with low and high phonemic rates, the location of the theta–alpha peak remains unchanged (low- and high-rate stimuli were selected to be those with phoneme rates lower than the 20^th^ percentile, or higher than the 80^th^ percentile, respectively. The stimulus file `audiobook_1` was excluded due to its high weight in the dataset [see Figure 1a]). In particular, the theta-alpha peak in the coherence spectrum is not moved to lower frequencies for the stimuli with low phonemic rate, and not moved to higher frequencies for stimuli with high phonemic rate, as would be expected if the peak was directly shaped by responses to the rate of phonemes. It is not clear whether the coherence magnitude itself is directly affected by phoneme rate, or instead some other acoustic difference between the speech material which contribute to the low- and high- phoneme rate groups.

**Figure 3:**
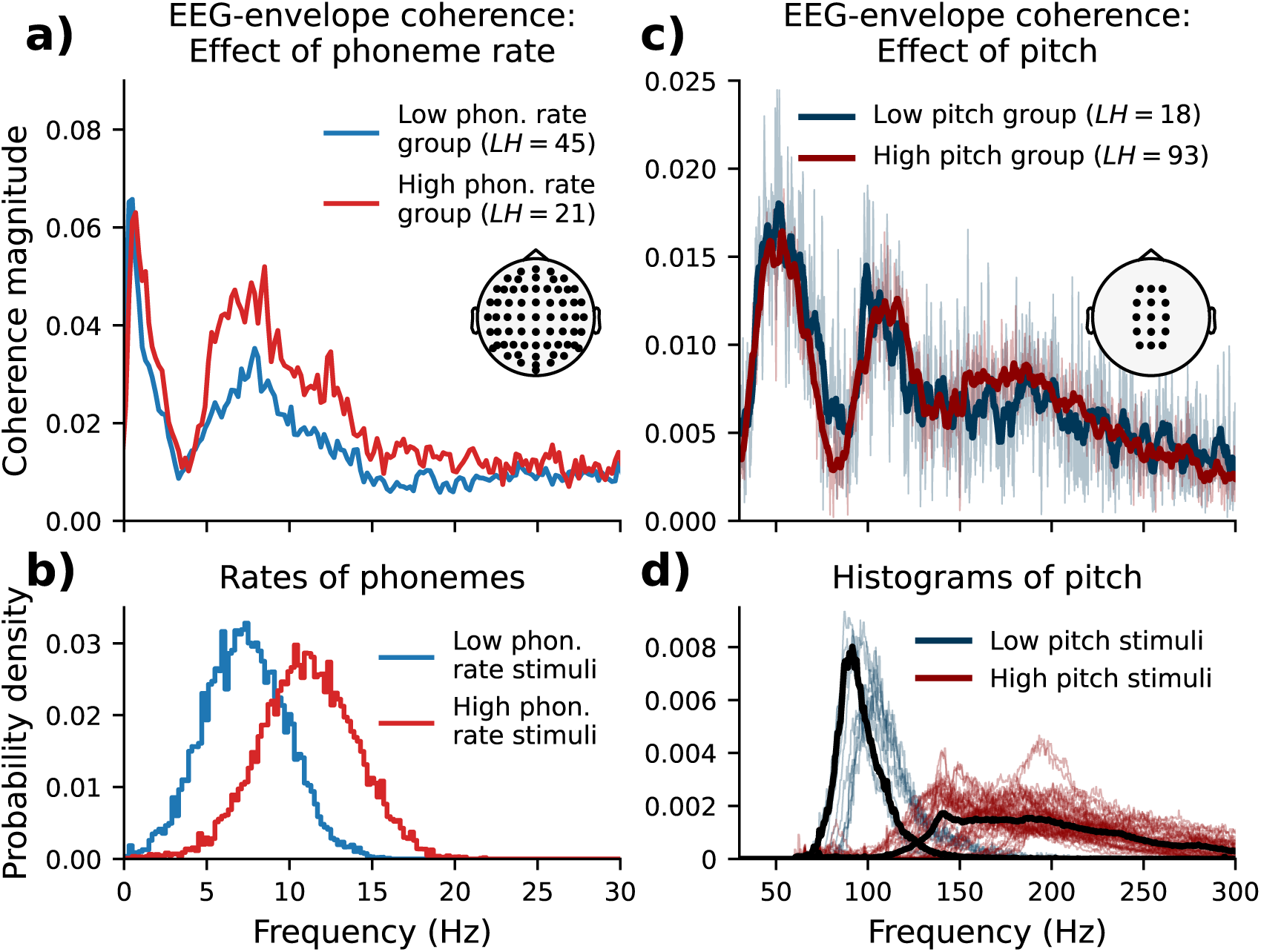
Testing the effect of phoneme rate and pitch on EEG–speech envelope spectral coherence. **a)** Root-mean-square coherence across all EEG channels for stimulus subgroups with lower and higher phoneme rates. The coherence profiles exhibit a similar overall shape, with no shift as a function of phoneme rate, although theta–alpha coherence is higher for the high-rate group. **b)** Weighted-average histograms of phoneme rates for the two groups. **c)** Equivalent analysis for low- and high-pitched stimuli. Thin lines show magnitude of coherence (which was averaged over a subset of central electrodes to improve SNR), and thick lines show the same data after smoothing with a 6 Hz boxcar window. The similarity of the profiles indicates that EEG-speech envelope coherence does not reflect pitch tracking. **d)** Histograms of pitch for individual stimuli (coloured lines) and group weighted averages (black).

#### Independence of speech pitch and the coherence spectrum

Because the SparrKULee dataset includes multiple talkers with different fundamental frequencies, the pitch distribution is bimodal, with maxima that roughly coincide with the peaks at 95 Hz and at 190 Hz in the coherence spectrum (Figure 2). To assess whether this alignment is coincidental, the speech material was divided into low- and high-pitched groups (selection criteria, respectively: third quartile of the F0 histogram *<* 130 Hz; first quartile *>* 130 Hz. The stimulus file `audiobook_1` was excluded due to its high weight in the dataset [see Figure 1a]). Figure 3c compares the coherence spectra for these groups alongside their corresponding pitch histograms. For visualisation, the complex coherence was averaged across a subset of central channels prior to computing magnitude, enhancing the detectability of the highest-frequency peak. The resulting coherence spectra for the two groups are remarkably similar, indicating that the peaks are not caused by the fundamental frequency and that high-gamma envelope tracking is largely invariant to speech pitch.

#### Coherence in the low-delta frequency range

Figure 2 shows that the low-delta peak exhibits the strongest coherence values. Unlike the other peaks, coherence in this range was strongly right-lateralised. The RMS coherence magnitude peaked at 0.6 Hz, consistent with the typical rate of intonational units. Because the speech material showed limited variability in prosodic structure, and because the intonation unit is just one of many possible measures of speech prosody, it was not possible to determine whether this peak indeed reflected of a neural mechanism for tracking prosody.

#### Dependence of EEG–envelope coherence on acoustic carrier band

The coherence spectrum shown in Figure 2 was derived from the broadband speech envelope, which was itself obtained by averaging across all channels of a gammatone cochleogram. To examine how envelope tracking depends on acoustic carrier frequency, spectral coherence was additionally computed between the EEG signals and each individual cochleogram channel.

The resulting sub-band coherence profiles are shown separately for low (0–30 Hz) and high (30–300 Hz) modulation frequencies (Figures 4a and 4b, respectively). In the low delta band (*<*2 Hz) as well as in the theta-alpha band (4–15 Hz), coherence was broadly distributed across cochleogram centre frequency. In contrast, coherence in the gamma range exhibited more pronounced carrier-frequency-dependent structure. Specifically, coherence in the 30–60 Hz and 80–120 Hz bands was strongest for cochlear channels with centre frequencies between approximately 500 Hz and 5 kHz, whereas coherence in the highest gamma band (140–220 Hz) was more narrowly confined to channels between roughly 2 and 5 kHz. These frequency ranges are closely aligned with those of the second and third formants (5th–95th percentiles: 705–3,345 Hz and 1,985–4,465 Hz, respectively). Estimates of the formant frequencies were tightly clustered within these ranges across all stimuli, precluding meaningful further analysis of whether carrier-band dependence adapted to formant frequency.

**Figure 4:**
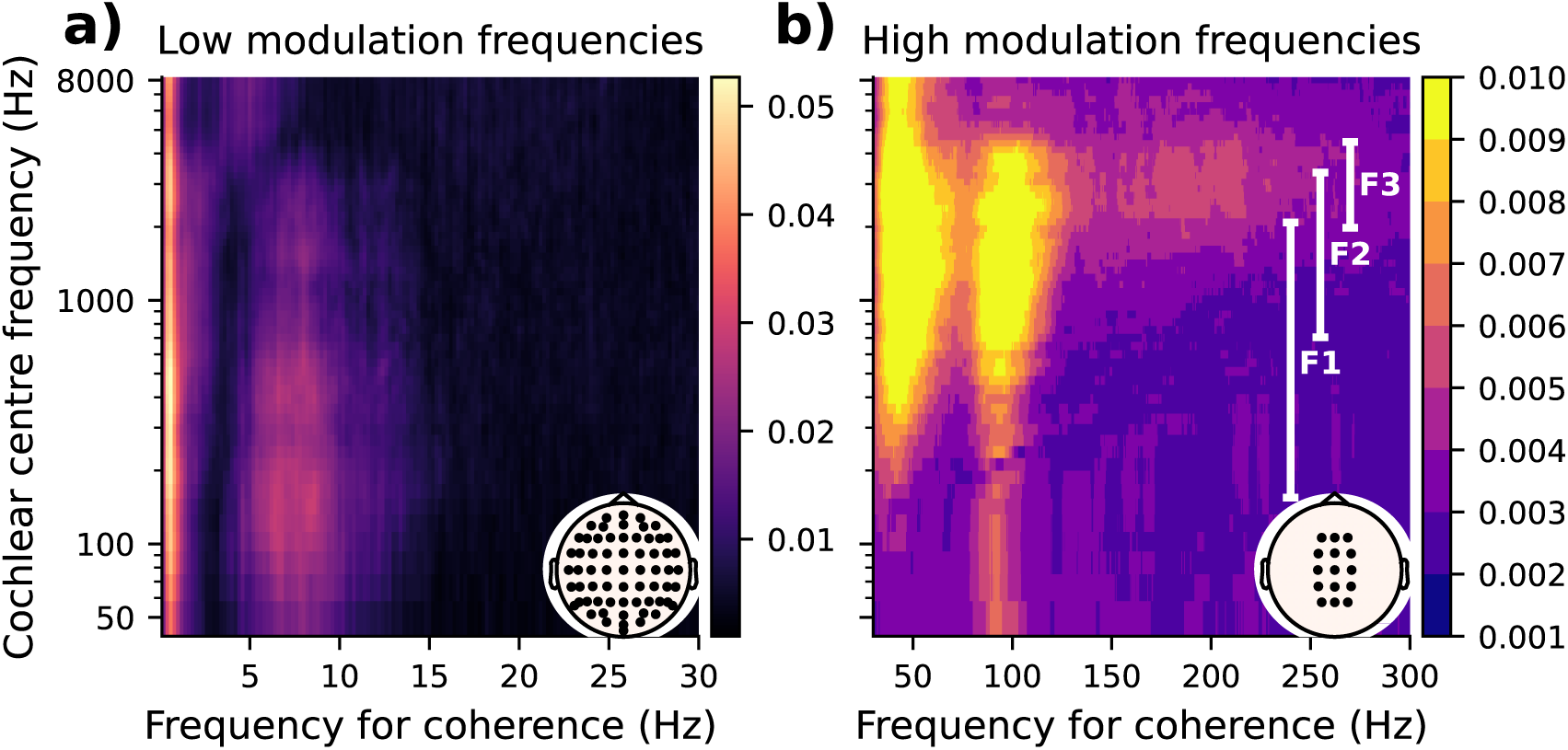
Coherence between EEG signals and individual cochleogram channels. **a)** Sub-band coherence at lower modulation frequencies (0–30 Hz) is broadly distributed across cochleogram channels with little fine structure. **b)** Sub-band coherence at higher modulation frequencies (30–300 Hz). In the ranges 30–60 Hz and 80–120 Hz, peak coherence occurs for cochleogram channels with centre frequencies between approximately 500 Hz and 5 kHz, whereas in the highest modulation band (140–250 Hz) it is more narrowly concentrated between 2 and 5 kHz. The 5th–95th percentiles of the first three formant frequency distributions are shown for comparison. Sensor legends in the bottom right of panels a) and b) indicate the EEG channels contributing to the RMS coherence estimates. Only the relevant channels identified in Figure 2a were included in panel b) to improve the signal-to-noise ratio of the sub-band coherence estimates.

### 3.2 Gamma-band responses to the speech envelope: generators and temporal profile

The coherence profile shown in Figure 2 exhibited a clear structure within the gamma range that was independent of speech pitch. Instead, neural tracking of the speech envelope was strongest within three distinct modulation bands: 30–60 Hz, 80–120 Hz, and 140–250 Hz. Clear differences in the spatial distribution of coherence between the highest modulation band and both lower bands are evident in the topographic plots of Figure 2, suggesting at least two distinct neural generators associated with these modulation-frequency ranges. To characterise the latencies of the responses in the different frequency bands, temporal response functions (TRFs) were fitted separately within each band.

#### Narrow-band TRFs in the gamma range

The TRFs for the three frequency bands are shown in Figure 5. Because these bands are relatively narrow, the resulting TRFs are oscillatory and extend broadly in time. To estimate the temporal locus of the underlying neural responses, we computed the Hilbert envelope of each TRF and measured its peak latency.

**Figure 5:**
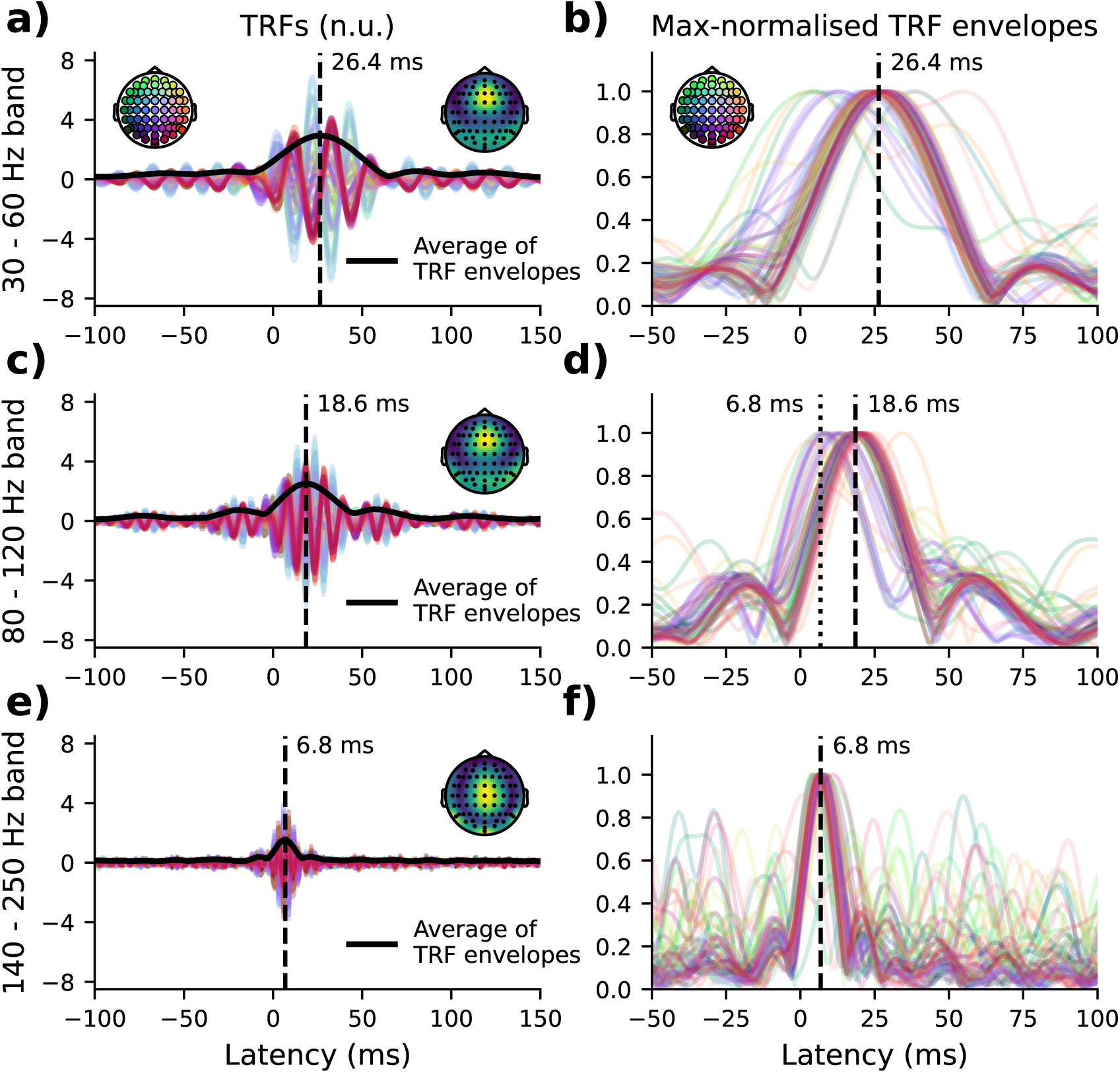
Temporal response functions (TRFs) and their Hilbert envelopes for the three narrow frequency bands 30–60 Hz (panels **a**–**b**), 80–120 Hz (**c**–**d**), and 140–250 Hz (**e**–**f**). The left column shows the TRFs in normalised units (n.u.), due to the EEG normalisation convention described in Section 2.7. Thick black lines indicate the mean Hilbert envelope across channels, and the accompanying topographic plots show the spatial distribution of TRF envelope values at the latency corresponding to the peak of this average envelope. The right column shows the channel-wise Hilbert envelopes of the TRFs. To facilitate comparison across channels, these envelopes were normalised by dividing each by its maximum value. In panels **b** and, more prominently, **d**, a subset of channel envelopes exhibit peaks at substantially earlier latencies than the channel-average envelope, with some as early as 6.8 ms. This pattern is consistent with the presence of an earlier neural generator that contributes more strongly at higher modulation frequencies.

Figure 5 shows that the narrow-band response latency decreased with increasing modulation frequency. Inspection of the channel-wise TRF envelopes indicated that this pattern arose from the superposition of two overlapping components with latencies of approximately 7 ms and 26 ms, with the relative contribution of the earlier component increasing at higher modulation frequencies. The 80–120 Hz band provided the clearest illustration of this structure, with a cluster of centroparietal channels peaking earlier than a second cluster of frontocentral channels. Comparison of the lowest (30–60 Hz) and highest (140–220 Hz) frequency bands further suggested that the contribution of the later component diminished as modulation frequency increased. Owing to the narrowband filtering, the two components were only partially separable in time.

To characterise the relative contributions of the two components as a function of modulation frequency, we fitted TRFs in 10 Hz-wide bands with centre frequencies ranging from 30 to 300 Hz (5 Hz spacing). The envelopes of the narrow-band TRFs are shown in Figure 6a, and their latencies in Figure 6b. The latencies decreased linearly between 30 and 150 Hz, consistent with a diminishing contribution of the later component at higher frequencies. The apparent response latency stabilises at a value of 6.8 ms above 150 Hz, indicating that only the early component contributed to the far-field EEG response in this range (Figure 6b). In conclusion, the choice of frequency band (including both width and centre frequency) strongly influences the interpretation of TRF latency in the gamma range.

**Figure 6:**
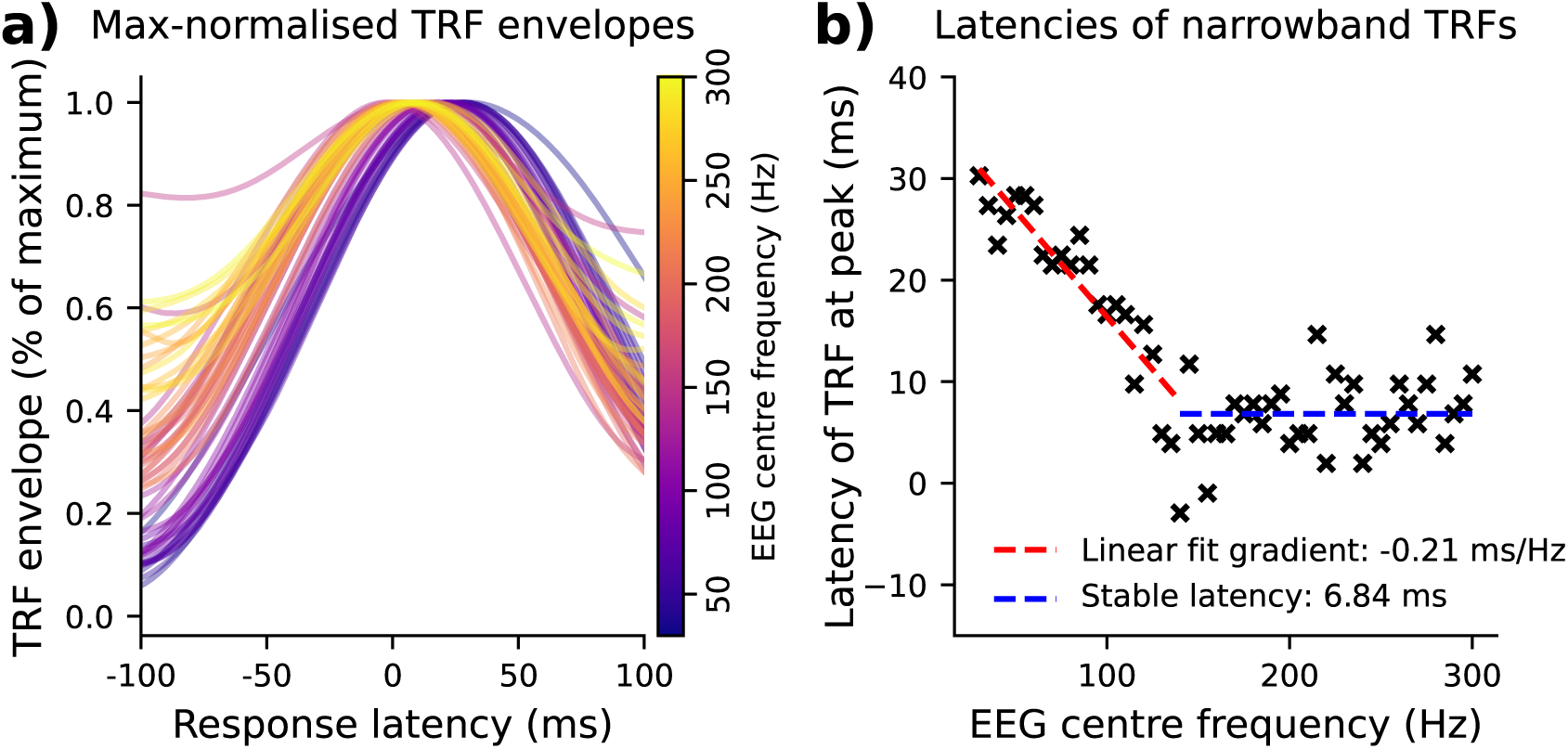
Temporal response functions (TRFs) estimated in narrow (10 Hz-wide) frequency bands. **a)** Channel-averaged Hilbert envelopes of the TRFs obtained for different centre frequencies. The averaged envelopes were each normalised through dividing by their global maxima. The latencies of the peaks decrease as the centre frequency increases. **b)** Latency of the TRF peak as a function of centre frequency. The early-latency component, with a latency of 6.8 ms, dominates at high centre frequencies above 150 Hz. Below 150 Hz, the apparent latency increases towards that of the late component, reflecting the stronger contribution from this component at lower frequencies.

#### Broadband TRFs in the gamma range

With this in mind, we additionally fitted broadband TRFs spanning 30–220 Hz (Figure 7). As anticipated, the increased bandwidth provided clearer temporal separation between the two components, with the TRF envelope resolving into two distinct peaks at delays of 6.8 ms and 23.4 ms, respectively. The shape of the broadband TRF should be interpreted in light of the modulation- and carrier-frequency dependence of the underlying sources. It likely also reflects far-field interference between the underlying neural generators, particularly at latencies between the temporal locii of the early and late contributions.

**Figure 7:**
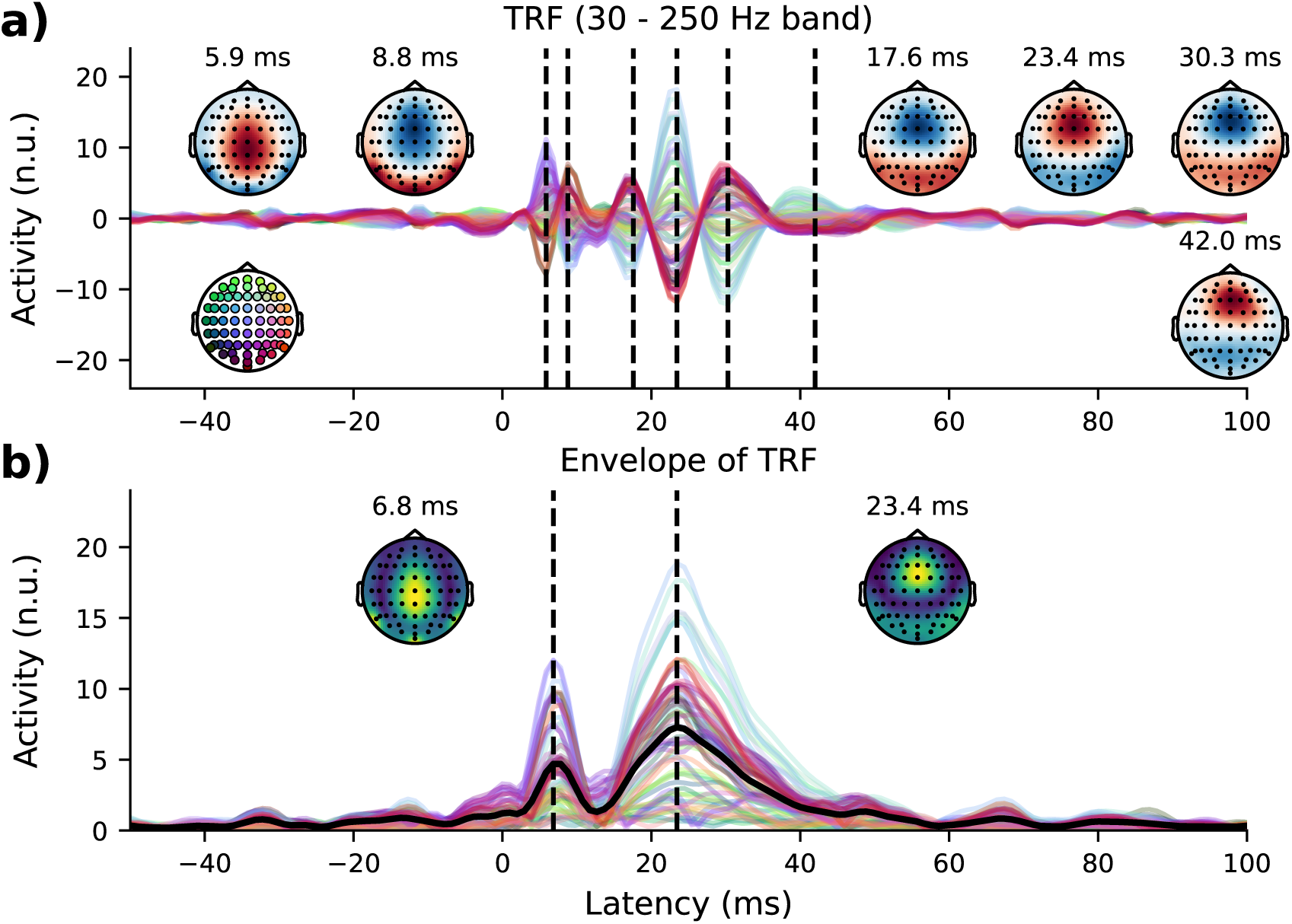
Temporal response functions (TRFs) estimated over the broad gamma band (30–250 Hz). **a)** Broadband TRF with scalp topographies shown at latencies exhibiting pronounced activity. **b)** Channel-wise Hilbert envelopes of the TRF, as well as channel average (black). The increased bandwidth improves temporal resolution, revealing two distinct peaks at 6.8 ms and 24.4 ms.

The TRFs shown in Figure 7 were derived using the channel-wise normalisation described in Section 2.7. Due to this normalisation, which effectively performed channel-wise division of the EEG signals by their standard deviation in the 30-250 Hz band, the TRFs did not directly capture the physical response strength of each EEG channel. Instead, they represented response strength as a fraction of total channel power (within the 30-250 Hz frequency band). Therefore, the TRFs of Figure 7 emphasised channels which were best explained by the linear signal model of equation 5, which is why the topographic plots resembled those of coherence as shown in Figure 2 (since coherence itself can be considered a narrow-band goodness-of-fit measure, as discussed in Section 2.8).

TRFs fitted in the 30-250 Hz frequency band without this normalisation are shown in Supplementary Figure S1. The latencies of those TRFs were unchanged from the latencies marked in Figure 7. However, the channels had a different scaling without this normalisation, which had an influence on the topographic maps inset into the TRF plots. Specifically, TRFs that were estimated without EEG normalisation exhibited a somewhat broader spatial profile, and the later aspects of the response displayed a right-hemispheric bias that was not present in the TRFs of Figure 7.

## 4 Discussion and conclusions

Neural envelope tracking is often linked to prosodic structure in the low-delta range, syllable-level processing in the theta band, and pitch periodicity in the high-gamma range. While a growing body of work supports neural speech processing at these timescales, our results show that far-field EEG measures of envelope tracking do not directly reflect syllabic processing or pitch tracking. In the following sections, we compare the spectral profile of EEG–speech envelope tracking with the rates of these features in turn, and discuss the potential mechanisms that could shape the observed coherence spectrum.

### 4.1 Neural tracking of prosody

The EEG–speech envelope coherence showed a peak in the low delta band, matching the rate of into-national units. The coherence in these low frequencies exhibited a clear right-hemisphere bias. Similar lateralisation has been reported in the MEG studies of Bourguignon et al. (2012) and Gross et al. (2013), with both demonstrating stronger responses for natural speech than for reversed or hummed speech. Bourguignon et al. (2012) further identified a common right-lateralised source for both understood and non-understood speech, alongside a distinct—though still right-lateralised—source for humming. Together, these findings suggest the presence of right-lateralised cortical networks that respond differentially to speech and non-speech sounds. Such low-frequency hemispheric asymmetries remain under-characterised, particularly compared with known theta and low-gamma asymmetries (see e.g. McGettigan and Scott, 2012; Poeppel, 2003).

Bourguignon et al. (2012) suggested that low-frequency envelope tracking might directly reflect prosodic processing, since the peak coherence frequency (0.5 Hz) roughly corresponds to the rate of prosodic units in natural speech. However, as demonstrated by the coincidental alignment between phoneme rate and the theta–alpha peak in our data, such correspondences do not necessarily imply a causal relationship. Future studies that manipulate prosodic structure could help determine whether low-frequency envelope tracking is specifically linked to prosodic processing.

### 4.2 Neural tracking of syllables, words and phonemes

Because the onsets of syllable nuclei coincide with sharp rises in the speech envelope, neural envelope tracking near the syllabic rate is often interpreted as reflecting a mechanism for syllable segmentation. Indeed, Oganian and Chang (2019) employed high-density cortical surface recordings to demonstrate specialised neural populations that respond to rising edges of the acoustic envelope at syllabic rates. Somewhat surprisingly, our results show that such neural syllable tracking is not visible in the coherence between the speech envelope and the EEG signals. Although neural envelope tracking was strongest in the 4–15 Hz frequency range, the syllabic rate in the SparrKULee stimuli occurred at a substantially slower frequency of approximately 2–5 Hz. Thus, theta-band envelope tracking—as measured in the far-field via scalp EEG—does not directly index syllable-level processing.

The absence of a word-rate peak in the coherence spectrum of Figure 2 suggests that the speech envelope is a weak acoustic marker of word boundaries. Indeed, many M/EEG studies of continuous speech perception explicitly model word onsets derived from time-aligned transcriptions, and show that these predictors explain neural variance beyond that captured by the speech envelope alone (Broderick et al., 2018; Weissbart et al., 2020). Furthermore, Ding et al. (2015) employed an elegant paradigm to demonstrate neural tracking of higher-level linguistic structure at the rate of words. In the context of these findings, we can conclude that despite strong evidence for the existence of word-level neural speech processes, these are not reflected well by envelope-based measures of neural tracking in far-field EEG measurements.

Figure 2 exhibited an alignment between the rate of phonemes and the theta-alpha peak between 4-15 Hz. However, subsequent analysis showed that this alignment was coincidental rather than reflective of actual phoneme tracking: speech material with slower and faster phonemic rates resulted in the same coherence profile, with no difference in the peak locations. Instead, studies that do report robust neural tracking of phonemes typically model categorical or expectancy-related features—such as phoneme surprisal or entropy—rather than relying on envelope-derived proxies for phoneme boundaries (Brodbeck et al., 2018; Di Liberto et al., 2015).

### 4.3 EEG-speech envelope coherence in the gamma band

The spectral coherence profile in the gamma band exhibited three peaks at approximately 45, 95, and 190 Hz. Many studies of gamma-band neural envelope tracking assume enhanced responses near the talker’s fundamental frequency, based on the hypothesis that envelope fluctuations associated with pitch periodicity in voiced speech drive strong neural responses. These studies typically employ band-limited analyses that exclude contributions from the low-gamma range. A notable exception is the study by Shan and Maddox (2025), which also reported coherence maxima corresponding to the two lowest gamma-band peaks reported here. Importantly, we found that even large variations in talker pitch produced little change in the gamma-band coherence profile, indicating that high-gamma envelope tracking in far-field EEG did not primarily reflect the pitch periodicity of continuous speech. The weaker contribution near 190 Hz has not previously been characterised in studies of continuous speech perception.

Using simple periodic stimuli, Tichko and Skoe (2017) characterised the amplitude of the EEG frequency-following response (FFR) as a function of stimulus frequency, producing a profile that closely resembles the EEG–speech envelope coherence spectrum reported here. Their results were obtained by averaging EEG responses across repeated presentations of triangle-wave stimuli. Because those stimuli were presented with a constant polarity (i.e. without polarity alternation), the resulting FFRs primarily track the amplitude fluctuations of these harmonically rich stimuli (Aiken & Picton, 2008). Therefore, the similarity between the results of that study and those reported here likely reflects common neural sources involved in encoding the acoustic amplitude envelope.

### 4.4 Neural sources of gamma-band envelope tracking

Our TRF analysis indicated that gamma-band envelope tracking was dominated by two neural generators with latencies of approximately 7 ms and 25 ms. The earlier component is highly consistent with wave V of the click-evoked auditory brainstem response, as reported in prior studies on continuous speech (Bachmann et al., 2021; Maddox & Lee, 2018). Polonenko and Maddox (2021) showed that far-field activity from the early generator—presumably located in the rostral brainstem—can be isolated by high-pass filtering neural responses above 150 Hz. Our results are consistent with this observation and further indicate that this source exhibits large-scale neural synchrony at modulation rates between 140 and 250 Hz. Our analysis of the narrow-band TRFs also indicated that the early generator contributed at lower gamma-band modulation rates. However, its modulation-frequency dependence below ∼120 Hz could not be resolved, as the later component dominated the far-field EEG response in the lower-gamma regime.

Polonenko and Maddox (2021) noted that the later component of the gamma-band TRF is consistent with wave Pa of the click-evoked middle-latency response (MLR), which has itself been associated with multiple stages along the thalamocortical pathway (Musiek & Nagle, 2018). Consistent with this, our findings suggest that both thalamic and cortical generators contribute to the later aspects of gamma-band neural envelope tracking.

The spatial pattern of coherence in the 30–60 Hz and 80–120 Hz bands exhibits a focal fronto-central maximum, closely matching the topographies of the later TRF components shown in Figure 7. Because both spectral coherence, as well as the TRFs obtained from normalised EEG, emphasise the reliability of envelope tracking relative to overall signal power (see Section 2.7), these data highlight generators that exhibit particularly faithful neural envelope tracking in the lower gamma range, independent of physical response magnitude. The observed topographies are consistent with a source dipole oriented predominantly within the sagittal plane; a radially oriented dipole located within the thalamus would be compatible with this pattern. This interpretation is further supported by the fact that, at higher stimulus rates, subcortical structures exhibit more robust neural phase locking than cortical sources (Joris et al., 2004).

At the same time, our analysis indicates that cortical contributions are not entirely absent in EEG measurements of gamma-band neural envelope tracking. When TRFs were estimated without channel-wise normalisation (Supplementary Figure S1), a right-hemispheric bias emerged at longer latencies (∼30 ms), consistent with the findings of prior MEG work which preferentially captured cortical contributions to envelope tracking (Kulasingham et al., 2020; Riegel et al., 2024; Schüller et al., 2024). This component is less prominent in the normalised analysis and does not strongly influence the coherence results, likely reflecting weaker large-scale phase locking in cortical populations compared with subcortical structures. Together, these findings suggest that gamma-band envelope tracking in far-field EEG is dominated by subcortical activity, with additional cortical contributions that are less robustly captured by coherence-based measures.

### 4.5 Carrier-band dependence of neural envelope tracking

The carrier-band analysis (Figure 4) revealed a clear difference in the carrier-frequency dependence between the two sources that contributed to gamma-band neural envelope tracking. The earlier source responded primarily to amplitude modulations in the 2–5 kHz carrier range (at least within the 140–250 Hz modulation band), whereas the later source showed a broader contribution of carrier frequencies from approximately 500 Hz to 5 kHz. The upper limits of these ranges may have been influenced by the frequency response of the ER-3A insert earphones, which was approximated during preprocessing using a low-pass filter with a 3.5 kHz passband edge and a broad transition extending to about 5.5 kHz. Despite some methodological differences, Polonenko and Maddox (2021) reported a broadly similar pattern, with carrier bands below 2 kHz producing substantially weaker wave-V-like TRFs. This contrasts with the click-evoked ABR literature, which robustly demonstrates that wave V can be elicited across a wide range of cochlear frequency bands (Abdala & Folsom, 1995; Don & Eggermont, 1978). The absence of a comparable contribution of carrier frequencies below 2 kHz in continuous-speech paradigms therefore remains unexplained, but may reflect limitations of the stimulus regressors (e.g. speech envelope or glottal pulse trains in (Polonenko & Maddox, 2021)).

In an MEG study, Hertrich et al. (2011) proposed that carrier-frequency dependence of neural envelope tracking may reflect the distribution of speech formants. Indeed, we observed an alignment between the typical ranges of the second and third formants and the carrier bands showing the strongest EEG–envelope coherence. From this perspective, the carrier-band dependence of EEG–speech envelope coherence may reflect the greater acoustic energy concentrated near formant frequencies. However, the comparatively weak contribution of the first formant remains unexplained. Further progress could be made using complex stimuli with systematic manipulations to the distribution of energy across carrier bands, or by employing TRF models with regressors that explicitly track formant dynamics.

### 4.6 Possible origins of the spectral coherence profile

Our results raise the question: if the structured spectral profile of EEG–speech envelope tracking does not primarily reflect the tracking of speech units or pitch, what mechanism gives rise to it? Plausible explanations include (i) intrinsic modulation-rate selectivity within the auditory system, (ii) resonances arising from endogenous oscillatory dynamics, and (iii) far-field interference between temporally-separated neural sources.

The first mechanism (i) assumes that distinct neural generators contribute preferentially at different modulation frequencies. Our data are consistent with cortical contributions that dominate at low frequencies below 30 Hz, midbrain structures contributing most strongly in the 30–120 Hz range, and earlier brainstem responses dominating at higher modulation frequencies above 150 Hz. Prior studies employing single-neuron recordings have demonstrated modulation-rate tuning within each of these anatomical regions. Although this tuning varies across neurons, its average across a population of neurons does play a role in shaping large-scale activity that is measurable non-invasively with EEG (Joris et al., 2004).

The second possible mechanism (ii) builds on the entrainment of intrinsic, large-scale oscillatory rhythms that are measurable with M/EEG. Such entrainment to speech rhythmicity has been proposed to support functions including temporal segmentation and feature binding (Meyer, 2017; Schroeder & Lakatos, 2009), and has been shown to contribute to neural speech tracking when measured with invasive electrophysiology (Akkol et al., 2025). If oscillatory mechanisms contribute substantially to the far-field EEG response, stimulation near the natural frequency of the underlying neural circuitry could lead to resonant amplification, manifesting as maxima in the coherence profile. A key prediction of oscillation-based accounts of speech processing is that theta-band activity becomes entrained to the syllabic rate, for example through phase resetting at acoustic edges associated with syllable nuclei onsets (Giraud & Poeppel, 2012; Meyer, 2017). However, the coherence between EEG and the speech envelope did not exhibit a corresponding peak at the syllabic rate. This suggests that, if oscillatory mechanisms contribute in the theta band, their influence is either weak in the EEG far-field, or masked by other processes that shape neural envelope tracking in this frequency range. A recent study also reported evidence for neural entrainment to speech at substantially higher frequencies near 100 Hz, at both cortical and sub-cortical levels of the auditory hierarchy (Coffey et al., 2021). Therefore, it is plausible that oscillatory mechanisms could give rise to the coherence maximum at 95 Hz reported in the present work.

The third mechanism (iii) was initially proposed by Galambos et al. (1981) to account for the local maximum in auditory steady-state response (ASSR) amplitude at a stimulus rate of 40 Hz. Modelling showed that the regular spacing between different waves of the middle-latency response, separated by delays of approximately 25 ms, could give rise to enhanced responses around 40 Hz under periodic stimulation. Separately, Tichko and Skoe (2017) proposed a model in which the relative latencies of six distinct neural generators caused far-field interference. By adjusting parameters such as neural source amplitude and relative source latency, they demonstrated that this model could reproduce EEG-measured frequency-following response (FFR) amplitudes as a function of stimulus fundamental frequency. In particular, the modelled responses included broad peaks at about 40 Hz, 80 Hz and 200 Hz. Because these are comparable to the peaks in the gamma range that we observe in the EEG coherence, the interference mechanism is likely play a large role in shaping the coherence spectrum that was observed in the gamma band.

### 4.7 Conclusions

During continuous speech listening, neural populations exhibit large-scale synchronisation to the speech envelope across a wide frequency range, from the low delta band starting at around 0.5 Hz to the high gamma band extending to about 200 Hz. The peaks emerging in the coherence, however, largely do not reflect speech structures. Notably, the maxima were not aligned with prominent speech features such as syllabic rate or pitch periodicity. The frequency dependence of the coherence appears to arise instead from far-field interferences of multiple neural generators and, potentially, oscillatory mechanisms.

These findings have important implications for the interpretation of neural envelope tracking. In particular, band-limited analyses should be approached with caution: our results show that the choice of frequency band can strongly influence both the apparent strength and latency of the response. Selecting bands based on the temporal structure of speech (e.g. syllabic rate) may therefore be misleading, while narrowband filtering can distort the temporal profile of the response and obscure the contribution of temporally distinct neural components.

## Data and Code Availability

We analysed a large, publicly-available EEG dataset (Accou et al., 2024).

## Author Contributions

M. T. performed the research, analyzed the data, interpreted the results and wrote the paper. T. R. interpreted the results and edited the paper.

## Funding

This project was supported by the German Federal Ministry of Research, Technology and Space (Cluster4Future, SEMECO, project number 03ZU1210FB), as well as by the German Science Foundation (DFG, project numbers 523344822 and 576735417).

## Declaration of Competing Interests

The authors declare that they have no competing interests.

## Supplementary Material

Supplementary Material (created during production as a web link to online material).

## Notes

### Competing Interest Statement

The authors have declared no competing interest.

## References

1. Abdala, C., & Folsom, R. C. (1995). Frequency contribution to the click-evoked auditory brain-stem response in human adults and infants. The Journal of the Acoustical Society of America, 97 (4), 2394–2404. 10.1121/1.411961

2. Accou, B., Bollens, L., Gillis, M., Verheijen, W., Van hamme, H., & Francart, T. (2024). Sparrkulee: A speech-evoked auditory response repository from KU Leuven, containing the EEG of 85 participants. Data, 9 (8), 94. 10.3390/data9080094

3. Ahn, E., & Chodroff, E. (2022). VoxCommunis: A corpus for cross-linguistic phonetic analysis. Proceedings of the 13^th^ Language Resources and Evaluation Conference, 5286–5294. https://aclanthology.org/2022.lrec-1.566/

4. Aiken, S. J., & Picton, T. W. (2008). Envelope and spectral frequency-following responses to vowel sounds. Hearing Research, 245 (1–2), 35–47. 10.1016/j.heares.2008.08.004

5. Akkol, S., Mishra, A., Markowitz, N., Espinal, E., Keshishian, M., Mesgarani, N., Schroeder, C., Mehta, A. D., & Bickel, S. (2025). Neural entrainment by speech in human auditory cortex revealed by intracranial recordings. Progress in Neurobiology, 253, 102823. 10.1016/j.pneurobio.2025.102823

6. Arai, T., & Greenberg, S. (1998). Speech intelligibility in the presence of cross-channel spectral asynchrony. Proceedings of the 1998 IEEE International Conference on Acoustics, Speech and Signal Processing, 2, 933–936. 10.1109/ICASSP.1998.675419

7. Bachmann, F. L., MacDonald, E. N., & Hjortkjær, J. (2021). Neural measures of pitch processing in eeg responses to running speech. Frontiers in Neuroscience, 15, 738408. 10.3389/fnins.2021.738408

8. Bain, M., Huh, J., Han, T., & Zisserman, A. (2023). Whisperx: Time-accurate speech transcription of long-form audio. Proceedings of INTERSPEECH 2023, 4489–4493. 10.21437/interspeech.2023-78

9. Biesmans, W., Das, N., Francart, T., & Bertrand, A. (2017). Auditory-inspired speech envelope extraction methods for improved EEG-based auditory attention detection in a cocktail party scenario. IEEE Transactions on Neural Systems and Rehabilitation Engineering, 25 (5), 402–412. 10.1109/TNSRE.2016.2571900

10. Boersma, P., & Weenink, D. (2021, January). Praat: Doing phonetics by computer [computer program, version 6.1.38]. https://www.praat.org

11. Bourguignon, M., De Tiège, X., de Beeck, M. O., Ligot, N., Paquier, P., Van Bogaert, P., Goldman, S., Hari, R., & Jousmäki, V. (2012). The pace of prosodic phrasing couples the listener’s cortex to the reader’s voice. Human Brain Mapping, 34 (2), 314–326. 10.1002/hbm.21442

12. Brodbeck, C., Hong, L. E., & Simon, J. Z. (2018). Rapid transformation from auditory to linguistic representations of continuous speech. Current Biology, 28 (24), 3976–3983. 10.1016/j.cub.2018.10.042

13. Broderick, M. P., Anderson, A. J., Di Liberto, G. M., Crosse, M. J., & Lalor, E. C. (2018). Electrophysiological correlates of semantic dissimilarity reflect the comprehension of natural, narrative speech. Current Biology, 28 (5), 803–809. 10.1016/j.cub.2018.01.080

14. Coffey, E. B. J., Arseneau-Bruneau, I., Zhang, X., Baillet, S., & Zatorre, R. J. (2021). Oscillatory entrainment of the frequency-following response in auditory cortical and subcortical structures. The Journal of Neuroscience, 41 (18), 4073–4087. 10.1523/jneurosci.2313-20.2021

15. Commuri, V., Kulasingham, J. P., & Simon, J. Z. (2023). Cortical responses time-locked to continuous speech in the high-gamma band depend on selective attention. Frontiers in Neuroscience, 17, 1264453. 10.3389/fnins.2023.1264453

16. Crosse, M. J., Di Liberto, G. M., Bednar, A., & Lalor, E. C. (2016). The multivariate temporal response function (mTRF) toolbox: A MATLAB toolbox for relating neural signals to continuous stimuli. Frontiers in Human Neuroscience, 10, 604. 10.3389/fnhum.2016.00604

17. de Jong, N. H., & Wempe, T. (2009). Praat script to detect syllable nuclei and measure speech rate automatically. Behavior Research Methods, 41 (2), 385–390. 10.3758/brm.41.2.385

18. Di Liberto, G. M., O’Sullivan, J. A., & Lalor, E. C. (2015). Low-frequency cortical entrainment to speech reflects phoneme-level processing. Current Biology, 25 (19), 2457–2465. 10.1016/j.cub.2015.08.030

19. Ding, N., Melloni, L., Zhang, H., Tian, X., & Poeppel, D. (2015). Cortical tracking of hierarchical linguistic structures in connected speech. Nature Neuroscience, 19 (1), 158–164. 10.1038/nn.4186

20. Ding, N., & Simon, J. Z. (2012). Emergence of neural encoding of auditory objects while listening to competing speakers. Proceedings of the National Academy of Sciences, 109 (29), 11854–11859. 10.1073/pnas.1205381109

21. Don, M., & Eggermont, J. J. (1978). Analysis of the click-evoked brainstem potentials in man using high-pass noise masking. The Journal of the Acoustical Society of America, 63 (4), 1084–1092. 10.1121/1.381816

22. Etard, O., & Reichenbach, T. (2019). Neural speech tracking in the theta and in the delta frequency band differentially encode clarity and comprehension of speech in noise. Journal of Neuroscience, 39 (29), 5750–5759. 10.1523/JNEUROSCI.1828-18.2019

23. Fontaine, B., Goodman, D. F. M., Benichoux, V., & Brette, R. (2011). Brian hears: Online auditory processing using vectorization over channels. Frontiers in Neuroinformatics, 5, 9. 10.3389/fninf.2011.00009

24. Forte, A. E., Etard, O., & Reichenbach, T. (2017). The human auditory brainstem response to running speech reveals a subcortical mechanism for selective attention. eLife, 6. 10.7554/elife.27203

25. Gabor, D. (1946). Theory of communication. Part 1: The analysis of information. Journal of the Institution of Electrical Engineers-part III: Radio and Communication Engineering, 93 (26), 429–441. 10.1049/ji-3-2.1946.0074

26. Galambos, R., Makeig, S., & Talmachoff, P. J. (1981). A 40-Hz auditory potential recorded from the human scalp. Proceedings of the National Academy of Sciences, 78 (4), 2643–2647. 10.1073/pnas.78.4.2643

27. Giraud, A.-L., & Poeppel, D. (2012). Cortical oscillations and speech processing: Emerging computational principles and operations. Nature Neuroscience, 15 (4), 511–517. 10.1038/nn.3063

28. Gross, J., Hoogenboom, N., Thut, G., Schyns, P., Panzeri, S., Belin, P., & Garrod, S. (2013). Speech rhythms and multiplexed oscillatory sensory coding in the human brain. PLoS Biology, 11 (12), e1001752. 10.1371/journal.pbio.1001752

29. Hertrich, I., Dietrich, S., Trouvain, J., Moos, A., & Ackermann, H. (2011). Magnetic brain activity phase-locked to the envelope, the syllable onsets, and the fundamental frequency of a perceived speech signal. Psychophysiology, 49 (3), 322–334. 10.1111/j.1469-8986.2011.01314.x

30. Inbar, M., Grossman, E., & Landau, A. N. (2025). A universal of speech timing: Intonation units form low-frequency rhythms. Proceedings of the National Academy of Sciences, 122 (34), e2425166122. 10.1073/pnas.2425166122

31. Issa, M. F., Khan, I., Ruzzoli, M., Molinaro, N., & Lizarazu, M. (2024). On the speech envelope in the cortical tracking of speech. NeuroImage, 297, 120675. 10.1016/j.neuroimage.2024.120675

32. Jadoul, Y., Thompson, B., & de Boer, B. (2018). Introducing Parselmouth: A Python interface to Praat. Journal of Phonetics, 71, 1–15. 10.1016/j.wocn.2018.07.001

33. Joris, P. X., Schreiner, C. E., & Rees, A. (2004). Neural processing of amplitude-modulated sounds. Physiological Reviews, 84 (2), 541–577. 10.1152/physrev.00029.2003

34. Keding, O., Alickovic, E., Skoglund, M. A., & Sandsten, M. (2024). Novel bias-reduced coherence measure for EEG-based speech tracking in listeners with hearing impairment. Frontiers in Neuroscience, 18, 1415397. 10.3389/fnins.2024.1415397

35. Kegler, M., Weissbart, H., & Reichenbach, T. (2022). The neural response at the fundamental frequency of speech is modulated by word-level acoustic and linguistic information. Frontiers in Neuroscience, 16, 2022.915744. 10.3389/fnins.2022.915744

36. Keitel, A., Gross, J., & Kayser, C. (2018). Perceptually relevant speech tracking in auditory and motor cortex reflects distinct linguistic features. PLoS biology, 16 (3), e2004473. 10.1371/journal.pbio.2004473

37. Kulasingham, J. P., Brodbeck, C., Presacco, A., Kuchinsky, S. E., Anderson, S., & Simon, J. Z. (2020). High gamma cortical processing of continuous speech in younger and older listeners. NeuroImage, 222, 117291. 10.1016/j.neuroimage.2020.117291

38. Lalor, E. C., Power, A. J., Reilly, R. B., & Foxe, J. J. (2009). Resolving precise temporal processing properties of the auditory system using continuous stimuli. Journal of Neurophysiology, 102 (1), 349–359. 10.1152/jn.90896.2008

39. Maddox, R. K., & Lee, A. K. C. (2018). Auditory brainstem responses to continuous natural speech in human listeners. eNeuro, 5 (1), e0441–17.2018. 10.1523/eneuro.0441-17.2018

40. McAuliffe, M., Socolof, M., Mihuc, S., Wagner, M., & Sonderegger, M. (2017). Montreal Forced Aligner: Trainable Text-Speech Alignment Using Kaldi. Proceedings of INTERSPEECH 2017, 498–502. 10.21437/Interspeech.2017-1386

41. McGettigan, C., & Scott, S. K. (2012). Cortical asymmetries in speech perception: What’s wrong, what’s right and what’s left? Trends in Cognitive Sciences, 16 (5), 269–276. 10.1016/j.tics.2012.04.006

42. Meyer, L. (2017). The neural oscillations of speech processing and language comprehension: State of the art and emerging mechanisms. European Journal of Neuroscience, 48 (7), 2609–2621. 10.1111/ejn.13748

43. Molinaro, N., & Lizarazu, M. (2018). Delta (but not theta)-band cortical entrainment involves speech-specific processing. European Journal of Neuroscience, 48 (7), 2642–2650. 10.1111/ejn.13811

44. Musiek, F., & Nagle, S. (2018). The middle latency response: A review of findings in various central nervous system lesions. Journal of the American Academy of Audiology, 29 (9), 855–867. 10.3766/jaaa.16141

45. Oganian, Y., & Chang, E. F. (2019). A speech envelope landmark for syllable encoding in human superior temporal gyrus. Science Advances, 5 (11), eaay6279. 10.1126/sciadv.aay6279

46. O’Sullivan, J. A., Power, A. J., Mesgarani, N., Rajaram, S., Foxe, J. J., Shinn-Cunningham, B. G., Slaney, M., Shamma, S. A., & Lalor, E. C. (2014). Attentional selection in a cocktail party environment can be decoded from single-trial EEG. Cerebral Cortex, 25 (7), 1697–1706. 10.1093/cercor/bht355

47. Peelle, J. E., Gross, J., & Davis, M. H. (2012). Phase-locked responses to speech in human auditory cortex are enhanced during comprehension. Cerebral Cortex, 23 (6), 1378–1387. 10.1093/cercor/bhs118

48. Poeppel, D. (2003). The analysis of speech in different temporal integration windows: Cerebral lateralization as ‘asymmetric sampling in time’. Speech Communication, 41 (1), 245–255. 10.1016/s0167-6393(02)00107-3

49. Polonenko, M. J., & Maddox, R. K. (2021). Exposing distinct subcortical components of the auditory brainstem response evoked by continuous naturalistic speech. eLife, 10, e62329. 10.7554/elife.62329

50. Radford, A., Kim, J. W., Xu, T., Brockman, G., McLeavey, C., & Sutskever, I. (2023). Robust speech recognition via large-scale weak supervision. Proceedings of the 40^th^ International Conference on Machine Learning, 28492–28518.

51. Riegel, J., Schüller, A., & Reichenbach, T. (2024). No evidence of musical training influencing the cortical contribution to the speech-frequency-following response and its modulation through selective attention. eNeuro, 11 (9), e0127–24.2024. 10.1523/eneuro.0127-24.2024

52. Rosen, S. (1992). Temporal information in speech: Acoustic, auditory and linguistic aspects. Philosophical Transactions of the Royal Society of London. Series B: Biological Sciences, 336 (1278), 367–373. 10.1098/rstb.1992.0070

53. Saberi, K., & Perrott, D. R. (1999). Cognitive restoration of reversed speech. Nature, 398 (6730), 760–760. 10.1038/19652

54. Saiz-Alía, M., & Reichenbach, T. (2020). Computational modeling of the auditory brainstem response to continuous speech. Journal of Neural Engineering, 17 (3), 036035. 10.1088/1741-2552/ab970d

55. Schroeder, C. E., & Lakatos, P. (2009). Low-frequency neuronal oscillations as instruments of sensory selection. Trends in Neurosciences, 32 (1), 9–18. 10.1016/j.tins.2008.09.012

56. Schüller, A., Schilling, A., Krauss, P., Rampp, S., & Reichenbach, T. (2023). Attentional modulation of the cortical contribution to the frequency-following response evoked by continuous speech. The Journal of Neuroscience, 43 (44), 7429–7440. 10.1523/jneurosci.1247-23.2023

57. Schüller, A., Schilling, A., Krauss, P., & Reichenbach, T. (2024). The early subcortical response at the fundamental frequency of speech is temporally separated from later cortical contributions. Journal of Cognitive Neuroscience, 36 (3), 475–491. 10.1162/jocna02103

58. Shan, T., & Maddox, R. K. (2025). Comparing methods for deriving the auditory brainstem response to continuous speech in human listeners. Imaging Neuroscience, 3, IMAG.a.19. 10.1162/imag.a.19

59. Tichko, P., & Skoe, E. (2017). Frequency-dependent fine structure in the frequency-following response: The byproduct of multiple generators. Hearing Research, 348, 1–15. 10.1016/j.heares.2017.01.014

60. Van Canneyt, J., Wouters, J., & Francart, T. (2021). Neural tracking of the fundamental frequency of the voice: The effect of voice characteristics. European Journal of Neuroscience, 53 (11), 3640–3653. 10.1111/ejn.15229

61. Vanthornhout, J., Decruy, L., Wouters, J., Simon, J. Z., & Francart, T. (2018). Speech intelligibility predicted from neural entrainment of the speech envelope. Journal of the Association for Research in Otolaryngology, 19 (2), 181–191. 10.1007/s10162-018-0654-z

62. Virtanen, P., Gommers, R., Oliphant, T. E., Haberland, M., Reddy, T., Cournapeau, D., Burovski, E., Peterson, P., Weckesser, W., Bright, J., van der Walt, S. J., Brett, M., Wilson, J., Millman, K. J., Mayorov, N., Nelson, A. R. J., Jones, E., Kern, R., Larson, E., . . . van Mulbregt, P. (2020). SciPy 1.0: Fundamental Algorithms for Scientific Computing in Python. Nature Methods, 17, 261–272. 10.1038/s41592-019-0686-2

63. Weissbart, H., Kandylaki, K. D., & Reichenbach, T. (2020). Cortical tracking of surprisal during continuous speech comprehension. Journal of Cognitive Neuroscience, 32 (1), 155–166. 10.1162/jocna01467

64. Zhang, Y., Zou, J., & Ding, N. (2023). Acoustic correlates of the syllabic rhythm of speech: Modulation spectrum or local features of the temporal envelope. Neuroscience & Biobehavioral Reviews, 147, 105111. 10.1016/j.neubiorev.2023.105111

